# NMY-2, TOE-2 and PIG-1 Regulate *C. elegans* Asymmetric Cell Divisions

**DOI:** 10.1101/2022.11.17.516947

**Authors:** Joseph Robinson, Jerome Teuliere, Shinja Yoo, Gian Garriga

**Author notes:** Institut de Systématique, Evolution, Biodiversité (ISYEB), Muséum national d’Histoire naturelle, CNRS, Sorbonne Université, EPHE, Université des Antilles, Paris, France. Department of Physiology, University of California San Francisco, San Francisco, CA 94143.

## Abstract

Asymmetric cell division (ACD) is an important mechanism that generates cellular diversity during development. Not only do asymmetric cell divisions produce daughter cells of different fates, many can produce daughters of different sizes, which we refer to as Daughter Cell Size Asymmetry (DCSA). In *C. elegans*, apoptotic cells are frequently produced by asymmetric divisions that exhibit DCSA, where the smaller daughter dies. We focus here on the divisions of the Q.a and Q.p neuroblasts, which produce apoptotic cells and divide with opposite polarity using both distinct and overlapping mechanisms. The PIG-1/MELK and TOE-2 proteins both regulate DCSA and specify the apoptotic cell fate in both the Q.a and Q.p divisions. In many asymmetric cell divisions, the non-muscle myosin NMY-2 is involved in properly positioning the cleavage furrow to produce daughters of unequal size. It was previously reported that NMY-2 is asymmetrically distributed and required for the DCSA of Q.a but not Q.p. In this study, we examined endogenously tagged reporters of NMY-2, TOE-2, and PIG-1 and found that all were asymmetric at the cortex during both the Q.a and Q.p divisions. TOE-2 and NMY-2 were biased toward the side of the dividing cell that would produce the smaller daughter, whereas PIG-1 was biased toward the side that would produce the larger daughter. We used temperature-sensitive *nmy-2* mutants to determine the role of *nmy-2* in these divisions and found that these mutants only displayed DCSA defects in the Q.p division. We generated double mutant combinations between the *nmy-2* mutations and mutations in *toe-2* and *pig-1*. The *nmy-2* mutations did not significantly alter the DCSA of the *toe-2* and *pig-1* mutants but did alter the fate of the Q.a and Q.p daughters. This finding suggests that NMY-2 functions together with TOE-2 and PIG-1 to regulate DCSA but plays an independent role in specifying the fate of the Q.a and Q.p descendants.

## Introduction

A core aspect of development is that a single cell can give rise to multiple cell types. This is often accomplished by ACD, where a cell divides to produce daughters with distinct fates. One mechanism contributing to ACD is the asymmetric distribution of cell fate determinants that specify daughter cell fates. Another mechanism contributing to ACD is DCSA, which results in daughter cells of unequal size. Different mechanisms can contribute to a shifted furrow that results in DCSA.

The *C. elegans* Q.a and Q.p neuroblast divisions provide examples of different mechanisms of DCSA (Fig 1). These sister cells both divide to produce a smaller daughter cell that dies while exhibiting opposite polarity: Q.a produces the smaller anterior Q.aa daughter, whereas Q.p produces the smaller posterior Q.pp daughter. Mutations that disrupt the size asymmetry also disrupt the apoptotic fate of the smaller daughter cell. Different DCSA mechanisms have been reported for the two cells: Q.a divides by a spindle-independent, HAM-1 and myosin-dependent mechanism, whereas Q.p divides by a spindle-dependent, HAM-1 and myosin-independent mechanism (1–3). Despite these differences, specific proteins are required for both divisions including the PAR-4/PIG-1 kinase pathway and the DEP-containing protein TOE-2 (4,5).

**Fig 1.**
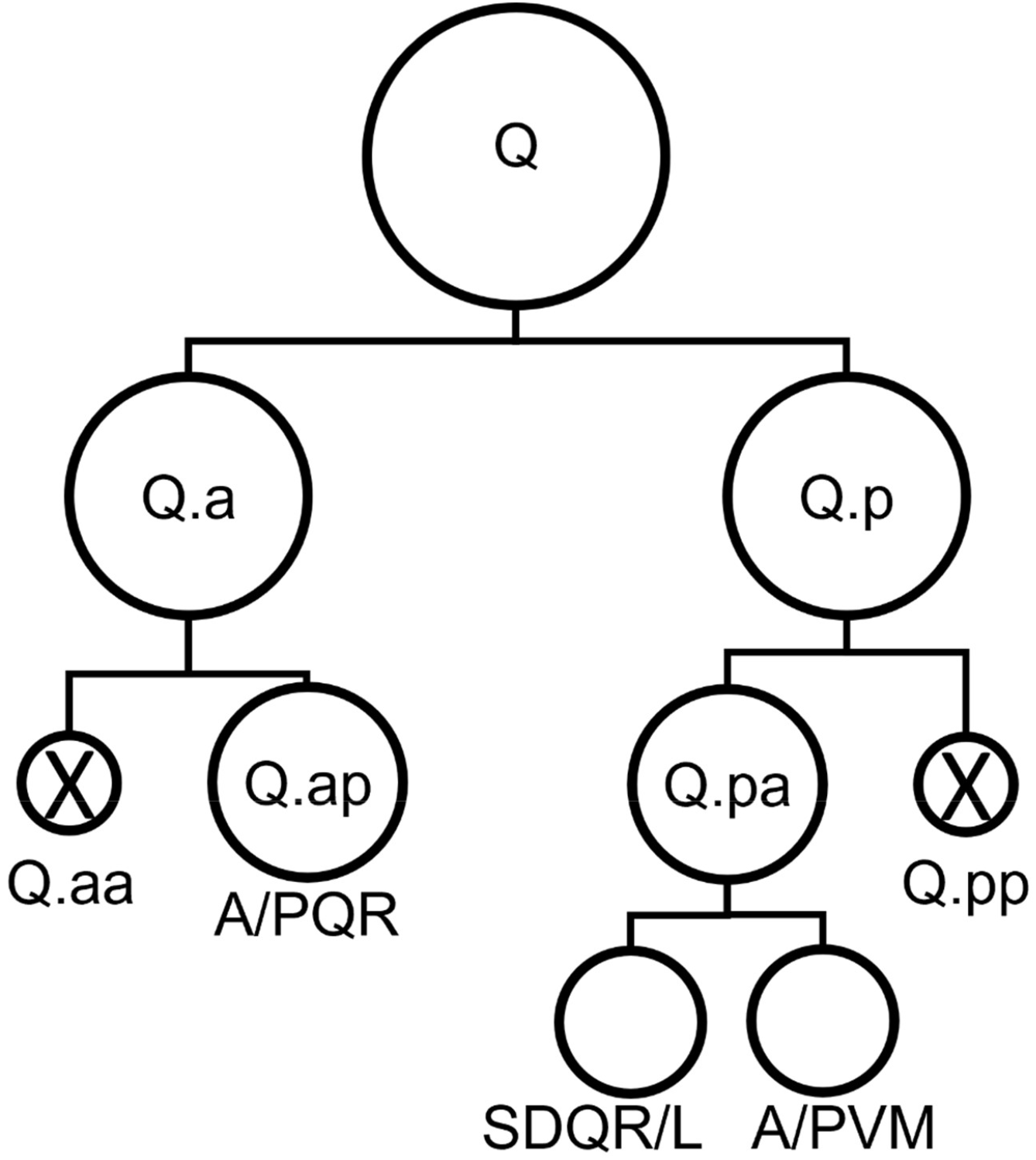
The *C. elegans* Q neuroblast divisions. The Q neuroblast divides to produce Q.a and Q.p, which divide to produce anterior (Q.aa or Q.pa) and posterior (Q.ap or Q.pp) daughter cells. In the Q.a division, the anterior daughter, Q.aa, is smaller and apoptotic. In the Q.p division, the posterior daughter, Q.pp, is smaller and apoptotic.

PIG-1 (*<u>p</u>ar-<u>1</u>*-like <u>g</u>ene) is the ortholog of the mammalian <u>M</u>aternal <u>E</u>mbryonic <u>L</u>eucine-zipper <u>K</u>inase (MELK) gene and a member of the AMPK family of kinases. In vertebrates, MELK has been shown to play a role in a wide range of cellular processes, including cell division, differentiation, death, and survival (6). In *C. elegans*, PIG-1 is involved in numerous asymmetric cell divisions, including the first embryonic division, the NSM neuroblast division, and both the Q.a and Q.p divisions (7,8). In several asymmetric cell divisions, PIG-1 has been shown to function downstream of a PAR-4/LKB1, STRD-1/STRADα, MOP-25.1,2/MO25α complex, presumably through phosphorylation of PIG-1 by PAR-4 kinase (4,7,9,10).

TOE-2 was originally identified by bioinformatics and shown biochemically to be a <u>T</u>arget <u>o</u>f <u>E</u>rk and was subsequently found to regulate both Q.a and Q.p ACD (5,11). TOE-2 has a DEP domain and multiple MPK docking sites (5,11). The <u>D</u>isheveled <u>E</u>gl-10 <u>P</u>leckstrin (DEP) domain is a protein-protein interaction domain typically found in transducers of cellular signaling near the cortex, particularly G-protein signaling. Using the predicted structure of TOE-2 from AlphaFold in an NCBI VAST search, we found that TOE-2 also has a region with structural similarity to RhoGAP domains (12–14). The structure of TOE-2 is similar to that of LET-99, a protein that also contains both a DEP domain and a degenerate RhoGAP-like domain and is involved in the first cell and EMS divisions (5,15,16). Based on the homology of the DEP domain and its predicted GAP domain, the mammalian homolog of TOE-2 is DEPDC7.

Non-muscle myosin is a central component of the actomyosin network and is involved in a wide variety of processes. The non-muscle myosin NMY-2 plays a key role in establishing the polarity of the first cell division and of the NSM, Q.a and Q.p neuroblast divisions (2,9,17). An interesting complication in generating a unifying model for NMY-2’s role in DCSA is that in these divisions, it localizes to different parts of the progenitor cell relative to the cell’s polarity. In the Q.a division, NMY-2 localizes to the anterior side, which will become the smaller daughter cell fated to die (2). This is similar to its pattern in Drosophila neuroblasts (18). In the NSM neuroblast, however, NMY-2 localizes to the side that will become the larger surviving NSM daughter cell. This is similar to the *C. elegans* first cell division where NMY-2 initially localizes to the side that will produce the larger AB blast cell (19). However, in Q.p, NMY-2 does not appear to be asymmetric (2). Despite the differences in NMY-2 asymmetry, PIG-1 regulates DCSA in all of these *C. elegans* divisions, and has been shown to regulate NMY-2 localization in the NSM and first cell divisions (4,7,9). Considering that in most of these divisions NMY-2 has been shown to play a role in furrow positioning independent of spindle positioning, the fact that the localization of NMY-2’s is different in these asymmetric divisions is puzzling.

We used endogenously tagged reporters of NMY-2, TOE-2, and PIG-1 to determine their subcellular localization during the Q.a and Q.p cell divisions in order to better characterize their function. We found that all were asymmetric at the cortex in both the Q.a and Q.p divisions, with TOE-2 and NMY-2 being biased towards the side of the progenitor that will produce the apoptotic daughter, while PIG-1 was biased towards the other side. We used temperature-sensitive *nmy-2* mutants to determine the role of *nmy-2* in these divisions and found that these alleles had only mild effects on the DCSA of the Q.p division and none on the Q.a division. When combined with mutations in *toe-2* or *pig-1*, the *nmy-2* alleles did not significantly alter the Q.a or Q.p DCSA defects of the *toe-2* or *pig-1* mutants but did alter the fate of their daughters. These findings suggest that NMY-2 plays a DCSA-independent role in specifying the fate of the Q.a and Q.p daughter cells.

## Materials and Methods

### Strains and Genetics

General handling and culture of nematodes were performed as previously described (20). N2 Bristol was the wild-type strain, and experiments were performed at 20°C unless otherwise noted.

The following mutations, integrated arrays and endogenously tagged genes were used:

LG I. *nmy-2(ne1490ts, ne3409ts)* (21), *nmy-2(cp13)* (*nmy-2::gfp+*LoxP) (22)

LG II. *toe-2(gm408ok2807)* (5), toe-2(syb1240) (mNeonGreen::*toe-2*) (this study)

LG III. *rdvIs1 [egl-17p::mCherry:his-24 + egl-17p::myristolated mCherry + pRF4]* (2), *casIs165[egl-17p::myr-mCherry; egl-17p::mCherry-TEV-S]* (23).

LG IV. *pig-1(gm280, gm301, gm344)* (8), *pig-1(syb2355)* (*pig-1::mNeonGreen*) (this study) LGX. *gmIs81 [mec-4p::mCherry, flp-12p::EBFP2, gcy-32p::gfp, egl-17p::gfp]* (5)

### Cell Count Protocol

Worms with the *gmIs81* integrated array were grown on Nematode Growth Media (NGM) seeded with OP50 at 15°C until the plates were populated with gravid adult hermaphrodites. Embryos were then collected after incubating the adults in.75 mL of a solution containing 500mM NaOH and 15% bleach until the adults were mostly dissolved, centrifuged to pellet the embryos which were then washed three times in 1.5mL of M9, and then plated on standard NGM plates seeded with OP50 and then transferred to 25°C. After 2-3 days, adult and fourth larval stage (L4) worms were transferred to 3-5 ul of 20mM Sodium Azide in M9 buffer on a 2% agarose pad. Hermaphrodites were scored for the number of observed Q lineage descendants. The number of observed PQR, SDQL and PVM neurons were scored in hermaphrodites with their left side up. The number of observed SDQR and AVM neurons were scored in hermaphrodites with their right side up. The numbers of all five cells were scored in hermaphrodites lying on their dorsal or ventral sides.

### Cell Count Analysis

The frequency of extra or missing cells was calculated for each cell type by dividing the number of sides with an extra or missing cell by the number of sides scored. AQRs were excluded from the final analysis as they were difficult to distinguish from other neurons in the head that express GFP in *gmIs81* animals.

The analysis of Q lineage division defects was predicated on the principle that certain cell-fate changes produce unique patterns that cannot be produced by a Q.aa or Q.pp cell that is normally fated to die surviving and adopting the fate of its sister cell or niece. For instance, occasionally Q.p will adopt the fate of Q.a leading to a missing A/PQR and extra SDQL/R or A/PVM cells. Another possibility is a progenitor may fail to divide, leading to the loss of its descendants. To eliminate lineages with these types of defects, each pattern of potential cell counts was analyzed to determine whether the production of extra neurons could result solely from a failure in apoptosis and transformation into its sister cell or niece. Only those lineages were counted in the filtered Q.a and Q.p results. The criteria for excluding a pattern were if a cell was missing or if there were three or more of any cell type. These patterns were arranged into defect categories corresponding with which division had failed. In the final analysis, we used two defect categories, QL.a and QL.p, with the patterns and categories described in Table

1. The frequency of each category was determined by the sum of worms exhibiting that category of defect divided by the number of worms scored for that lineage. Specifically, the frequency of QL.p defects was calculated as (# QL.p defective)/(# QL.p defective + # QL.p normal), while the frequency of QL.a defects was calculated as (# QL.a defective)/(# QL.a defective+ # QL.a normal).

### Imaging

Worms with the *gmIs81* integrated array were grown on NGM seeded with OP50 at 15°C until the plates had a large number of embryos and larvae. The plates were then put at 25°C for 4 hours. Worms were then washed off the plates with M9 and transferred to microcentrifuge tubes. They were then spun in a tabletop centrifuge for less than 6 seconds to pellet the larger, adult worms. The supernatant was then transferred using a glass pipette to a fresh microcentrifuge tube. This was then spun for 30 seconds and the supernatant was removed until only ∼100μL remained. 2 μL of 1M Sodium azide was added, and the tube was briefly vortexed and spun for 30 seconds. All but ∼10 μL of supernatant was removed, and, using a glass pipette, the pellet and remaining M9 + sodium azide were transferred to a 2% agarose slide. To determine the sizes of the daughter cells and the number of Q-derived neurons, the worms were imaged using an Axio Observer Z1 microscope.

For time-lapse imaging, worms were prepared as previously described (5). Time-lapse images of Q neuroblast divisions were captured with seven plane Z-stacks (Z-step: 0.5 µm) in 30-second intervals on a spinning-disk (CSU-X1; Yokogawa) confocal microscope. Images were captured using an EM CCD camera (Evolve; Photometrics) and SlideBook software (Intelligent Imaging Innovations).

### Image Analysis

Image analysis was performed using the FIJI package for ImageJ (24). To measure DCSA, the outlines of recently divided Q.p or Q.a daughters were traced using the lasso select tool in ImageJ, and the area was measured for each cell to determine the ratio. To measure the localization of the endogenously tagged reporters, we identified time points for metaphase, anaphase, telophase and cytokinesis in the time-lapse images and created sum Z-projections of the slices containing the best cross-section of the dividing cell. We then performed line scans around the cortex, using the segmented line and plot profile tools in FIJI, measuring the intensity of the endogenously tagged protein in the 488 nm channel and myristoylated mCherry in the 561 nm channel across 3 pixels every 1/6 microns. A line was drawn through either the metaphase plate or cleavage furrow and measurements started at a point where that line intersected the membrane and followed the entire cell cortex as marked by the myristoylated mCherry. The other point intersecting the line of the metaphase plate or cleavage furrow was then marked as were the anterior and posterior sides of the furrow. We also established the background levels of fluorescence by measuring the average intensity in each channel of a section of the body cavity that expressed neither reporter.

### Line-scan Modelling

To model the line-scan data, each point was normalized and paired with information about the relevant variables. First, the background intensity for each channel was subtracted from the intensity values. To normalize protein to membrane levels we divided the intensity at 488 nm by the intensity at 561 nm for each point, which we refer to as the Normalized Intensity Ratio. To compare cells of different sizes, we determined the Normalized Distance by dividing the distance from the start of the measurement by the total distance measured. We also determined the normalized distance between each point and the nearest metaphase plate or cleavage furrow point. Each point was then paired with its Normalized Intensity Ratio, the cell type, the phase of the division, and whether it was anterior or posterior to the division plane.

The line-scan information was used to construct Generalized Mixed Linear Models (GLMMs) using the MCMCglmm package in R (25). We used the Normalized Intensity Ratio as the dependent variable, the specific measurement as a random variable, and the Cell Type, the Phase, Anterior vs Posterior, and the distance from the furrow as fixed variables. We also added the interaction terms of Phase::Anterior vs Posterior and Phase::Furrow Distance to account for changes in their effects in different phases. The resulting models allowed us to estimate the effect size of each variable.

In order to determine whether there was a significant difference between the Anterior and Posterior effects during each phase, we used the emmeans R package to calculate the estimated marginal means for the Phase and Anterior vs Posterior interaction term (26). Using the emmeans pairs function to perform pairwise comparisons between the different estimated effect sizes, we found the difference between the estimated effect sizes for each pair of interaction pairs of Phase and Posterior vs Anterior. The estimated difference between the Anterior and Posterior in each phase is then used to calculate a z-ratio and Tukey adjusted p value. The sign of the estimated difference indicates the direction, with positive values indicating Posterior and negative values indicating Anterior. Plots were made using the gather_emmeans_draws function in the tidybayes R package to generate draws from the marginal posterior distributions of the models and the ggplot2 R package to generate the plots from those draws (27,28).

### RNA interference (RNAi)

For *nmy-2* RNAi experiments, the *nmy-2* RNAi bacteria from the Ahringer Library (29), or L4440 control were grown overnight in LB with 50 μg/ml carbenicillin. 20uL of O/N culture was spread on NGM plates with 50 μg/ml carbenicillin and 1 mM IPTG and allowed to grow at 25°C for 48hrs. Embryos were collected by bleaching and plated on RNAi or L4440 control plates and cell counts were performed as described above.

## Results

### NMY-2, TOE-2, and PIG-1 are asymmetrically distributed during the divisions of Q.a and Q.p neuroblasts

The *C. elegans* Q lineage has been a model for studying of asymmetric cell division. The left and right Q cells each undergo a series of divisions along the anterior-posterior axis (A-P) to generate three neurons and two cells fated to die (30). The Q daughters, Q.a and Q.p, each divide to generate daughter cells that are asymmetric in size and fate (Fig 1). Q.a divides to generate a smaller anterior daughter cell that dies and a larger posterior daughter that survives and differentiates into an A/PQR oxygen-sensing neuron. Q.p divides with the opposite polarity to generate a larger anterior daughter that survives and divides to generate the A/PVM mechanosensory neuron and the SDQR/L interneuron. Because the right and left Q and their descendants migrate in opposite directions, the neurons on the right side (AQR, AVM and SDQR) are in positions anterior to those on the left side (PQR, PVM and SDQL).

The non-muscle myosin NMY-2, the AMPK-family kinase PIG-1, and the DEPDC7 homolog TOE-2 all regulate the asymmetry of the Q.a and Q.p divisions (2,5,8). To better understand the roles of these molecules in asymmetric cell division, we took time-lapse images of the Q.a and Q.p divisions in strains containing endogenously tagged NMY-2::GFP, PIG-1::mNeonGreen, or mNeonGreen::TOE-2 (Fig 2). We then measured the intensity of the GFP or mNeonGreen at the cortex and normalized it to an mCherry cortical marker. Taking the average normalized intensity of the posterior and anterior sides of the cells, we observed a slight asymmetry of NMY-2::GFP (Fig 2A) and mNeonGreen::TOE-2 (Fig 2B) towards the side of Q.a or Q.p that will produce the daughter cell fated to die, and strong asymmetry of PIG-1::mNeonGreen towards the side that will produce the daughter cell that survives (Fig 2C).

**Fig 2.**
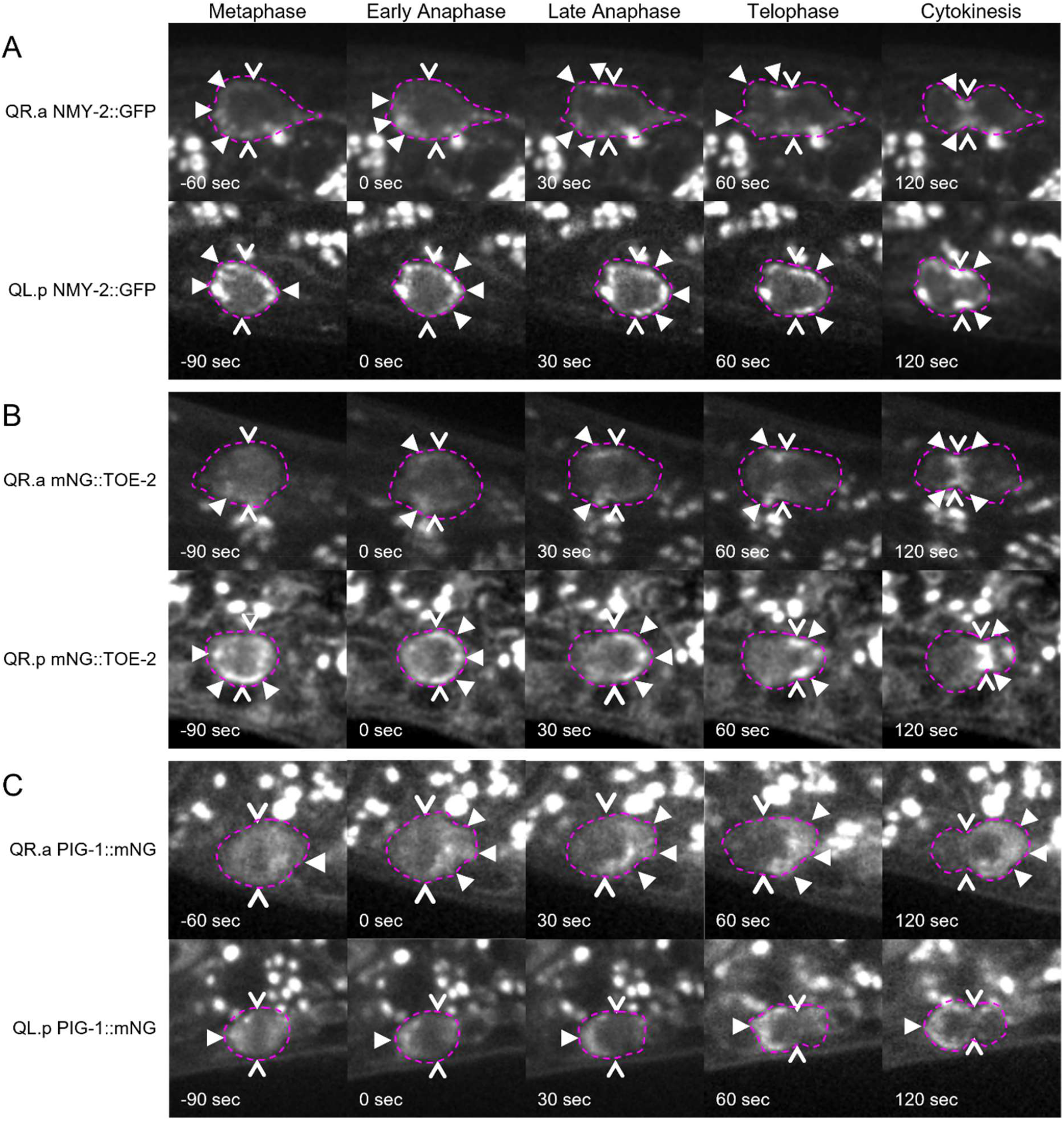
Time-lapse images of endogenously tagged DCSA proteins in the Q.a and Q.p divisions. A-C) Time-lapse imaging of QR.a and QL.p divisions in A) NMY-2::GFP, B) mNeonGreen::TOE-2, and C) PIG-1::mNeonGreen. Dotted lines outline the membrane of the cells. Filled arrowheads indicate areas of higher GFP or mNeonGreen signal, open arrowheads indicate the division plane.

Because both NMY-2 and TOE-2 localize to the cleavage furrow during anaphase and telophase, the ratio of the average intensity of the posterior and anterior could be misleading as identical levels of the protein flanking the furrow combined with the size asymmetry of the daughter cells would result in the smaller side having a higher average intensity. To mitigate this problem, we performed line scans around the cortex and annotated each measured point on the line scan with the intensity normalized by taking the ratio of the intensity of the GFP signal and the intensity of the mCherry cortical marker, the distance from the division plane normalized to the circumference of the cell, the phase of the division, and the positions anterior or posterior to the division plane.

We used this information to construct two GLMMs for each reporter, one for Q.a and one for Q.p, with the normalized intensity ratio as the dependent variable and the other information as fixed variables. We also included interaction terms between the phase and anterior or posterior as well as phase and furrow distance. Interaction terms account for the difference in the effect of one variable based on the value of another variable. In this model, the distribution of the protein with respect to both the A-P bias and proximity to the furrow can vary in different phases. The resulting models allowed us to estimate the effect size of each variable, and, most importantly, the interaction terms allowed us to compare the levels of the reporter at the anterior and posterior within each phase after accounting for the other variables. The resulting models’ estimates and parameters can be found in Table S1.

Using our model, we found that there was significantly more NMY-2::GFP at the anterior of Q.a during metaphase, anaphase, and telophase. During cytokinesis, NMY-2 was not asymmetric and localized to the cleavage furrow (Fig 3A). Our Q.p model had more NMY-2 at the anterior of Q.p at metaphase but no asymmetry during anaphase and telophase. NMY-2 localized asymmetrically to the posterior of Q.p during cytokinesis (Fig 3A).

**Fig 3.**
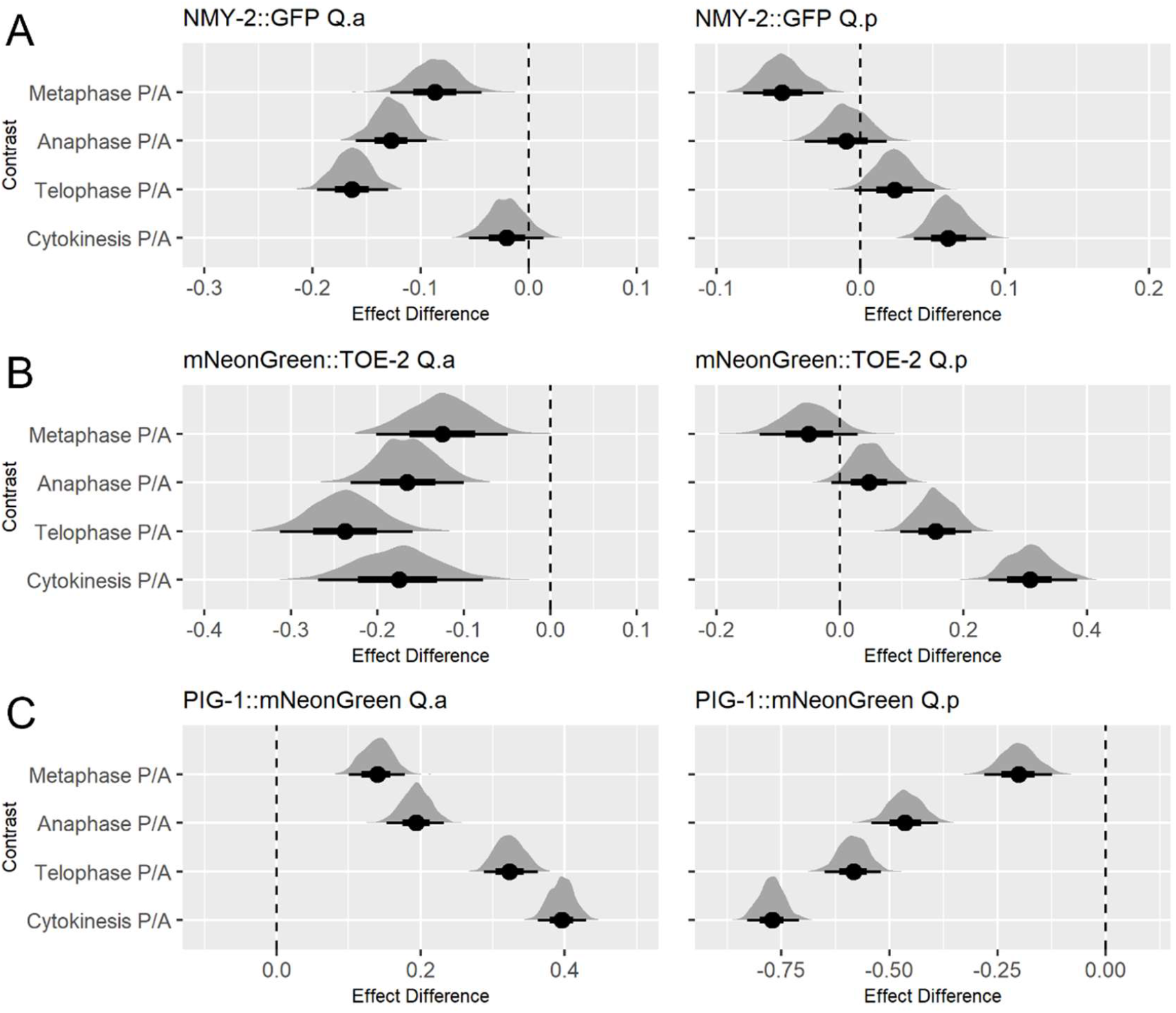
Line scan modelling analysis of endogenous reporters shows asymmetry in Q.a and Q.p divisions. A-C) Modelling analysis comparing anterior and posterior effect in Q.a and Q.p in Metaphase, Anaphase, Telophase and Cytokinesis for A) NMY-2::GFP, B) mNeonGreen::TOE-2, and C) PIG-1::mNeonGreen. The black bars represent the confidence intervals, while the distributions represent the frequency of draws of that value. Negative values indicate a greater anterior effect, positive values indicate a greater posterior effect.

Our model for the distribution of mNeonGreen::TOE-2 was similar to that of NMY-2::GFP: it had a greater anterior distribution throughout the Q.a division though, unlike NMY-2::GFP, mNeonGreen::TOE-2 remained asymmetric during cytokinesis (Fig 3B). During the Q.p division, TOE-2 moved steadily toward the posterior as the division progressed (Fig 3B). We also found that mNeonGreen::TOE-2 and NMY-2::GFP colocalized in the germline (Fig S1). NMY-2 localizes to the lateral membranes that separate at germline nuclei and accumulates at the ring channels that form the pores that connect the germ cells to a central canal, the rachis (31). mNeonGreen::TOE-2 localized to the junction between the germline progenitors Z1 and Z2 and continued to localize to the apical surface of the germline cells through all stages of development (Fig S2).

The distribution of TOE-2 reported here differed from previous descriptions of GFP-tagged transgenes. A GFP-tagged *toe-2* cDNA expressed from an *egl-17* promoter accumulated in the nuclei of interphase cells, but we detected no nuclear TOE-2 with the endogenously tagged gene. This is similar to the nuclear localization of the GFP-tagged Arf GEF GRP-1; the endogenous GRP-1 was not detectible with anti-GRP-1 antibodies (32). The nuclear localization of excess TOE-2 and GRP-1 might ensure that the proteins do not accumulate at the cortex in the interphase cells where they might have deleterious effects.

Our models showed that PIG-1::mNeonGreen was much more asymmetric than either NMY-2::GFP or mNeonGreen::TOE-2, localizing to the posterior of Q.a and the anterior of Q.p during mitosis. For both Q.a and Q.p, the PIG-1::mNeonGreen asymmetry increased as the cells progressed through mitosis (Fig 3C).

### Temperature-Sensitive *nmy-2* mutants reveal a role in Q lineage ACD

Although the asymmetry of NMY-2 in Q.a has been documented (2), we were surprised to find that its distribution was also asymmetric in Q.p during metaphase and cytokinesis. To determine whether NMY-2 also functioned in the Q.p division, we asked whether the two temperature-sensitive *nmy-2* mutants, *nmy-2(ne1490ts)* and *nmy-2(ne3409ts)*, had altered Q.a and Q.p DCSA when shifted to 25°C four hours before imaging. We detected a significant decrease in Q.p DCSA in both *nmy-2(ne3409ts)* (P<0.01) and *nmy-2(ne1490ts)* mutants (P<0.05) compared to the control. Neither had a significant effect on Q.a DCSA. (Fig 4C, D).

**Fig 4.**
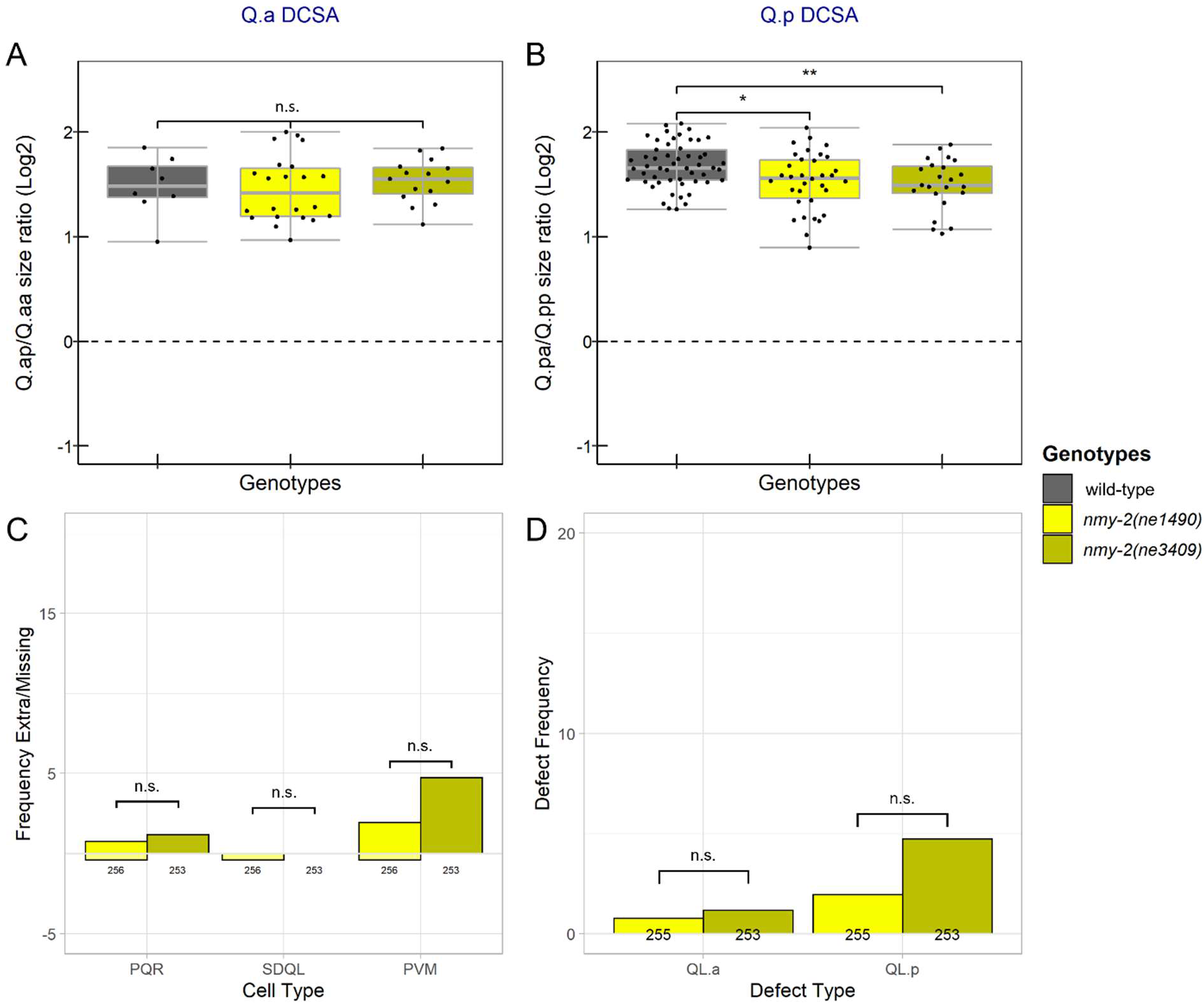
DCSA and cell-fate defects in temperature-sensitive *nmy-2* mutants. A,B) Box plots of the area ratios of A) Q.ap/Q.aa and B) Q.pa/Q.pp divisions in control and *nmy-2*(ts) mutants. C) Frequency of extra (positive y axis) and missing (negative y axis) Q lineage cells. D) Frequency of extra cell defects that could be explained by survival of QL.aa or QL.pp. (C, D) Below each bar are the number of lineages scored. *: P<0.05, **: P<0.01, ***: P<0.001, n.s.: P>0.05.

We also assessed the fates of the QL cell descendants in the mutants by counting the number of PQR, SDQL, and PVM neurons in adult hermaphrodites that had been shifted to 25°C during embryonic development. To determine the number of these cells, we used the *gmIs81* reporter, which labels each cell type with a different fluorescent marker (5). We did not count QR descendants because we could not reliably distinguish AQRs from other neurons expressing GFP in the head. The two *nmy-2* mutants had a low frequency of extra and missing QL descendants (Fig 4C). The wild-type control had no extra or missing cells (N=133).

Some of these phenotypes are difficult to interpret and may result from a failure of progenitor cells to divide or cell-fate transformations earlier in the lineage. For example, worms that lack Q.p descendants but have two or more Q.a descendants potentially represent Q.p to Q.a transformations. Because of these complications, we filtered the cell counts to remove all instances with either a missing cell or three or more of a given cell type. These filtered cell counts were then grouped based on whether they had QL.a or QL.p defects (Table 1). Both *nmy-2(ts)* alleles had increased frequencies of QL.a and QL.p defects compared to wild-type (Fig 4D). There was not a significant difference between the frequency of QL.a or QL.p defects between *nmy-2(ne1490ts)* and *nmy-2(ne3409ts)* raised at the nonpermissive temperature (Fig 4C,D).

**Table 1:** QL lineage Cell count defect filtering

| # Cells on Left Side |  |  | Defect Present? |  |
| --- | --- | --- | --- | --- |
| PQR | SDQL | PVM | QL.a | QL.p |
| 1 | 1 | 1 | No | No |
| 1 | 1 | 2 | No | Yes |
| 1 | 2 | 1 | No | Yes |
| 1 | 2 | 2 | No | Yes |
| 2 | 1 | 1 | Yes | No |
| 2 | 1 | 2 | Yes | Yes |
| 2 | 2 | 1 | Yes | Yes |
| 2 | 2 | 2 | Yes | Yes |

The filtering method used to determine what cell count patterns corresponded to each defect type for the left side Q lineage neurons while excluding all cases requiring a cell fate transformation beyond the survival of the QL.aa or QL.pp cell.

### Temperature-sensitive *nmy-2* alleles alter *toe-2* mutant Q lineage cell-fate defects but not Q.a or Q.p DCSA

The similar distributions of NMY-2 and TOE-2 during the Q.a and Q.p divisions suggest that they may function together to regulate DCSA. We constructed double mutants with the presumptive null *toe-2(gm408ok2807)* allele and each temperature-sensitive *nmy-2* allele to determine if the two genes function together or in parallel. Consistent with the two genes acting together, there were no significant DCSA differences between the *toe-2* single mutant and either *toe-2; nmy-2(ts)* double mutant (Fig 5 A,B). Because the *nmy-2* mutants do not alter Q.a DCSA, the lack of a *toe-2* enhancement is difficult to interpret, but because the mutants do alter Q.p DCSA, the lack of enhancement suggests that *nmy-2* and *toe-2* function together to regulate the size asymmetry of this division.

**Fig 5.**
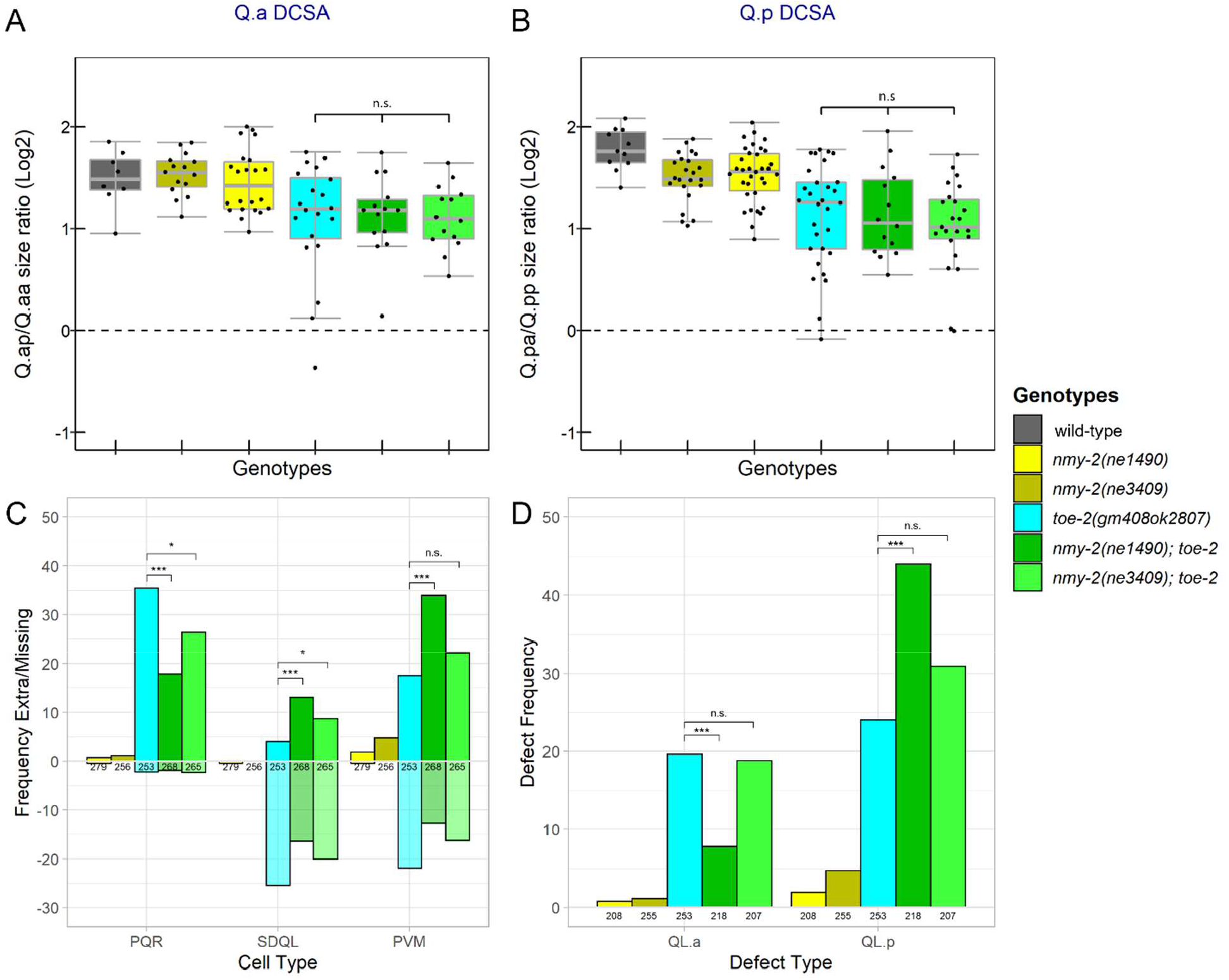
Temperature-sensitive *nmy-2* mutants enhance *toe-2* Q lineage cell fate but not DCSA defects. A, B) Box plots of area rations of A) Q.ap/Q.aa and B) Q.pa/Q.pp divisions. C) Frequency of extra (positive y axis) and missing (negative y axis) QL lineage cells. D) Frequency of extra cell defects that could be explained by survival of QL.aa or QL.pp with no other cell-fate transformations. (C, D) Below each bar are the number of lineages scored. *: P<0.05, **: P<0.01, ***: P<0.001, n.s.: P>0.05.

Our observations of cell fate in the single and double mutants were more complicated. When compared to *toe-2(gm408ok2807)*, the *nmy-2(ne1490ts); toe-2* strain had a significant (p<0.001) increase in the frequency of extra SDQL and PVM neurons and a decrease in the frequency of extra PQR neurons (Fig 5C). After filtering, there was a similar decrease in QL.a defects and an increase in QL.p defects in the double mutants (Fig 5 D). The *nmy-2(ne3409ts); toe-2* strain had weaker effects: a significant (P<0.05) increase in the frequency of extra SDQL cells and a decrease in the frequency of extra PQR cells (Fig 5C). However, the *nmy-2(ne3409ts); toe-2* strain was not significantly different from *toe-2(gm408ok2807)* in the frequency of QL.a or QL.p defects after filtering (Fig 5 D).

The difference between the results for the temperature-sensitive alleles led us to repeat the experiments with *nmy-2* RNA interference (RNAi) (Supplementary Table 1). We observed that *nmy-2* RNAi treatment of *toe-2(gm408ok2807)* resulted in an increase of both the Q.a and Q.p cell-fate defects relative to the *toe-2(gm408ok2807)* RNAi control. This observation is consistent with our Q.p findings for the *toe-2* double mutant with *nmy-2(ne1490ts)* allele. Unlike the suppression of the *toe-2* Q.a defects by *nmy-2(ne1490ts), nmy-2* RNAi enhanced the Q.a defects of *toe-2*. One explanation for this enhancement is RNAi of *nmy-2* is not specific and also reduces the function of the related gene *nmy-1*, but inconsistent with this possibility, *nmy-2* RNAi did not alter the levels of GFP in an *nmy-1::GFP* transgenic strain, while it did eliminate it in an *nmy-2::GFP* strain (not shown).

The lack of a significant enhancement of the *toe-2* mutant by the *nmy-2(ne3409ts)* mutation suggests that this allele is weaker than the *nmy-2(ne1490ts)* allele. To test this hypothesis, we treated the *nmy-2* mutants with *nmy-2 RNAi* and found both *nmy-2(ne1490ts)* and *nmy-2(ne3409ts)* resulted in a significant increase in the frequency of Q.p defects at the nonpermissive temperature, suggesting that both mutations reduce but do not eliminate *nmy-2* function at the nonpermissive temperature (Table S2). Because there was no effect on the *toe-2* Q.a defects by the *nmy-2(ne3409ts)* mutation and *nmy-2* RNAi enhanced these defects, the reason for the suppression of these defects by *nmy-2(ne1490ts)* is unclear. The *nmy-2(ne1490ts)* mutation could alter *nmy-2* function in a unique way or the strain could have a second mutation that leads to the suppression.

### Temperature-sensitive *nmy-2* alleles suppress *pig-1(gm301)* Q.a cell fate defects while not significantly altering Q.a DCSA

Our finding that PIG-1 and NMY-2 localized to opposite sides of Q.a suggests that these two molecules play different roles in these cells. To determine how *pig-1* and *nmy-2* interact, we constructed *nmy-2; pig-1(gm301)* double mutants and scored the single and double mutants for Q.a and Q.p DCSA defects and the presence or absence of Q-lineage neurons. There were no significant differences in DCSA of either Q.a or Q.p between *pig-1* single and *nmy-2(ts); pig-1* double mutants (Fig 6 A, B). Both double mutant strains had a significant decrease in the frequency of extra PQR neurons and QL.a-specific defects when raised at the nonpermissive temperature of 25°C. (Fig 6 C, D). The *pig-1* single and the *nmy-2; pig-1* double mutants displayed similar Q.p defects.

**Fig 6.**
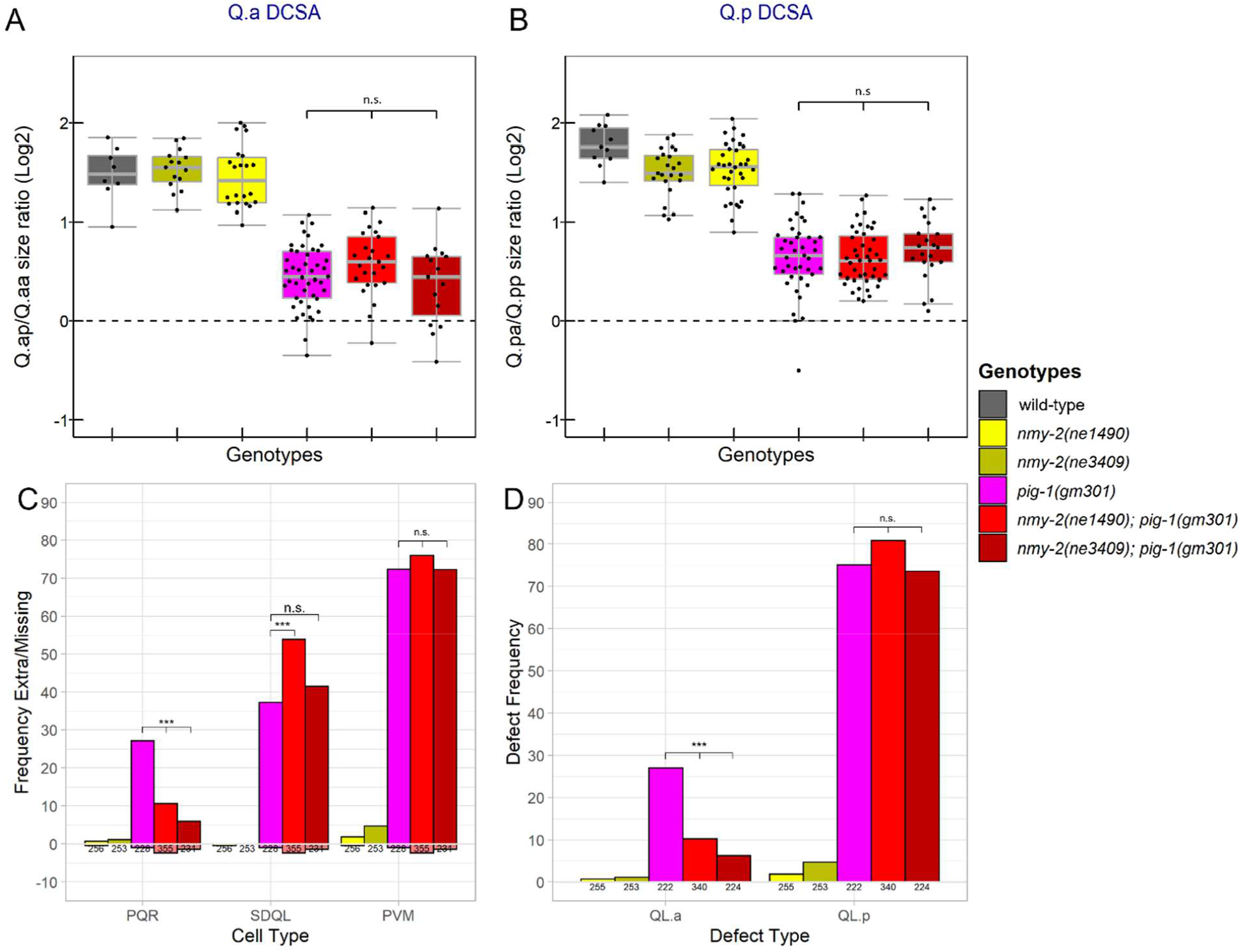
Temperature-sensitive *nmy-2* alleles suppress *pig-1(gm301)* Q.a cell fate defects while not significantly altering Q.a DCSA. A,B) Box plots of the area ratios of A) Q.ap/Q.aa and B) Q.pa/Q.pp divisions. C) Frequency of extra (positive y axis) and missing (negative y axis) QL lineage cells. D) Frequency of extra cell defects that could be explained by survival of QL.aa or QL.pp. (C, D) Below each bar are the number of lineages scored. *: P<0.05, **: P<0.01, ***: P<0.001, n.s.: P>0.05.

The significant increase in SDQLs in the *nmy-2(ne1490ts); pig-1(gm301)* strain compared to the *pig-1(gm301)* strain is potentially due to what is referred to by Mishra et. al (33) as an increase in mitotic potential (Fig 6C). Specifically, in instances where QL.pp survives in *pig-1(gm301)*, it divides 46.2% of the time, while in *nmy-2(ne1490); pig-1(gm301)* it divides significantly (P<0.001) more frequently, 63.8% of the time (Table S3).

### Time-lapse imaging of a *pig-1(gm344)* mutant background shows a reversal of NMY-2::GFP asymmetry in Q.a and Q.p

To further examine the potential interaction effects between *nmy-2* and *pig-1*, we constructed a *nmy-2::GFP; pig-1(gm344)* strain. We observed a reversal of the NMY-2::GFP asymmetry, with more posterior NMY-2::GFP in Q.a during metaphase, telophase and cytokinesis (Fig 7A,C), and a more anterior NMY-2::GFP in Q.p throughout mitosis (Fig 7 B,D). The Q.a reversal differs from a previous report that observed a loss of NMY-2::GFP asymmetry in a *pig-1* mutant (2).

**Fig 7.**
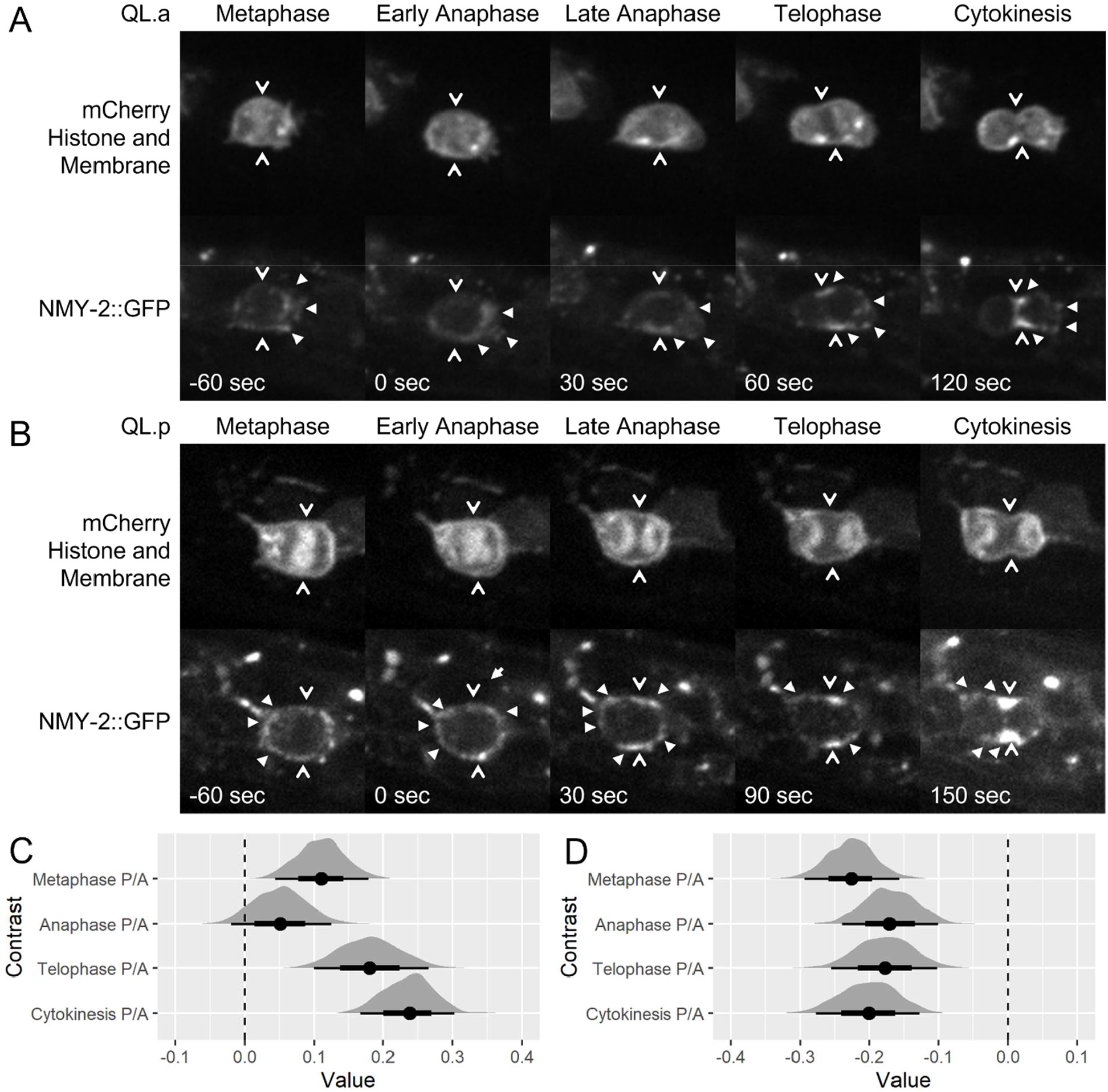
Time-lapse images of NMY-2::GFP in *pig-1(gm344)* background shows a reversal of NMY-2::GFP asymmetry in Q.a and Q.p. A) QR.a and B) QR.p live imaging of NMY-2::GFP in the *pig-1(gm344)* background. Filled arrows indicate areas of higher NMY-2::mNeonGreen signal, open arrows indicate the division plane. C, D) Line scan modelling analysis of C) Q.a and D) Q.p Posterior vs Anterior effect size difference. Negative values indicate a greater posterior effect, positive values indicate a greater anterior effect.

## Discussion

### NMY-2 is asymmetric in Q.a and Q.p and functions in Q.p DCSA

Our results for NMY-2 are interesting in that they do not align fully with previously reported findings. In particular, the experiments of Ou et al. using a GFP-tagged NMY-2 transgene showed significant asymmetry of the GFP::NMY-2 in Q.a but not Q.p (2). The authors also performed Chromophore Assisted Laser Inactivation (CALI) experiments where they inactivated the GFP::NMY-2 transgene at the anterior of Q.a or the posterior of Q.p during late anaphase (2). In Q.a, they saw that this perturbation caused an attenuation, loss, or reversal of DCSA and an increased rate of survival and differentiation of Q.aa. However, they observed no change in Q.p DCSA (2). This led them to conclude that *nmy-2* primarily regulates Q.a DCSA.

By contrast, we observed that endogenously tagged NMY-2::GFP exhibited asymmetry in both divisions, with a bias towards the anterior throughout the Q.a division and an increasing bias towards the posterior after an initial anterior bias in the Q.p division. Our experiments with temperature-sensitive *nmy-2* mutants showed no significant change in Q.a DCSA and a significant reduction in Q.p DCSA. Supporting this observation, we observed a higher frequency of extra Q.p lineage cells compared to Q.a.

What could explain these differences? Excess NMY-2 transgene expression could make it harder to observe the more subtle asymmetry we observed in Q.p imaging of the endogenous *nmy-2* tagged with GFP. The transgene used in the Ou et al. study contains multiple copies of *nmy-2* expressed from a heterologous promoter (34). A possible confounding factor in the CALI experiments is the presence of endogenous NMY-2. Whereas the CALI experiments would still produce an imbalance in contractile forces by reducing the levels of NMY-2 in the inactivated region, the expression of untagged NMY-2 makes it likely that NMY-2 activity or the activity of other nearby molecules affected by CALI still persists in the irradiated areas. The lack of a CALI result with irradiation of the posterior of Q.p could reflect less sensitivity to altered levels of NMY-2 than the anterior of Q.a.

Given the asymmetry of NMY-2 in Q.a and the results of CALI experiments reported by Ou et al., we were surprised to find no DCSA phenotype for Q.a in the *nmy-2* mutants. The lack of a Q.a DCSA phenotype could result from residual *nmy-2* function in the mutants: the low level of NMY-2 activity remaining would be sufficient to prevent loss of asymmetry in Q.a. In Drosophila neuroblasts, as in *C. elegans* Q.a and Q.p, more non-muscle myosin localizes to the side that will produce the smaller cell. In the Drosophila neuroblast, this localization is thought to produce a cortical contraction that prevents membrane extension (35). In the *C. elegans* NSM neuroblast and the first cell division, NMY-2 localizes to the opposite side of the dividing cell, the side that will produce the larger daughter cell (18). In the NSM neuroblast division, the *nmy-2(ne3409ts)* mutation results in a complete loss of DCSA. The authors proposed that NMY-2 creates cortical flows that are required to establish the gradient of cell-fate determinants and increase cortical contractility on the ventral side, which will produce the larger daughter cell (9). This model contrasts with the models of cortical contractility on the side that produces the smaller daughter cell. Further experiments are needed to determine which model would be correct in the context of the Q lineage.

### NMY-2 has a role in Q-lineage fate determination that is independent of its role in DCSA

The temperature-sensitive NMY-2 alleles caused defects in cell-fate determination in both Q.a and Q.p despite having no or minimal impact, respectively, on the DCSA of those divisions. These findings suggest that NMY-2 has a DCSA-independent role in cell fate determination in the Q.a and Q.p divisions. The finding that NMY-2 regulates cell fate has also been observed in the NSM neuroblast division. Besides eliminating DCSA, the *nmy-2(ne3409ts)* mutation disrupted the gradient of CES-1, a Snail-like transcription factor and cell-fate determinant (9).

We observed that the endogenously tagged mNeonGreen::TOE-2 and NMY-2::GFP reporters exhibited similar localization patterns in the Q.a and Q.p divisions, with both being biased towards the side that will produce the daughter cell fated to die as well as to the cleavage furrow. This localization suggests that they may have related functions. Consistent with this possibility, double mutants of *toe-2* and the temperature-sensitive alleles of *nmy-2* exhibit no increase in DCSA in either the Q.a or Q.p divisions when compared to *toe-2* on its own but did have differences in specifying the fate of their descendants. In particular, both *nmy-2(ts)* mutations increased the frequency of the *toe-2* mutant QL.p defects. The interactions between the *nmy-2* and *toe-2* mutations on the fate of the Q.a descendants were variable and difficult to interpret. The Q.p interaction is consistent with a role for NMY-2 in regulating cell fate independent of its role in DCSA. A DCSA-independent role is further supported by the finding that both *nmy-2(ts)* mutations suppressed the frequency of QL.a defects in *pig-1(gm301)* mutants without significantly altering QL DCSA.

A possible mechanism for NMY-2’s function in specifying fate in the Q lineage is to distribute cell fate determinants, as was shown for CES-1 in the NSM neuroblast (9). This has also been observed in Drosophila neuroblasts where non-muscle myosin is required for the basal distribution of two cell-fate determinants, Prospero and Numb (36). Another possible explanation would be an indirect effect wherein the *nmy-2(ts)* strains may have defects that slow cytokinesis or abscission, potentially resulting in cell fate determinants losing asymmetry due to a persisting connection between the daughter cells. This is supported by the observation of persistent intercellular bridges between the daughter cells of both Q.a and Q.p in strains containing the *nmy-2(ts)* alleles (Fig S3) as well as the fact that the original characterization of the *nmy-2(ts)* alleles reported that the cleavage furrow of the first embryonic division would halt or regress when the embryo was shifted to the non-permissive temperature (21).

### PIG-1 is asymmetric in Q.a and Q.p and regulates NMY-2 localization in those divisions

Previous studies using a *Ppig-1::pig-1::gfp* transgene showed that PIG-1::GFP localized to the cortex and centrosomes, but did not report any asymmetry in the Q lineage (4). The endogenously tagged PIG-1::mNeonGreen showed cortical localization as well as a clear asymmetry towards the side of the neuroblast that would produce the daughter cells fated to live during both the Q.a and Q.p divisions.

Using double mutant strains of *pig-1* and the temperature-sensitive *nmy-2* alleles, we observed no significant change in surviving QL.pp cells compared to *pig-1* on its own. We observed a shift of lineages that produced an extra SDQL and PVM to those that produced both extra SDQL and PVM and interpret this as an increase in mitotic potential. By contrast, there was a significant decrease in the frequency of surviving QL.aa cells in both *nmy-2(ne1490ts); pig-1* and *nmy-2(ne3409ts); pig-1* when compared to *pig-1* on its own.

We were surprised to observe a reversal of the NMY-2::GFP localization pattern in both Q.a and Q.p in the *pig-1* mutant background: using a transgene that expresses NMY-2::GFP, both Ou et al. and Wei et al. observed a loss of NMY-2 asymmetry in a *pig-1* mutant. It is noteworthy that there was no significant alteration in DCSA in either division for either *nmy-2; pig-1* double mutant strain when compared to the *pig-1* single mutant strain.

In the NSM neuroblast division, *nmy-2* functions downstream of *pig-1* (9). The authors showed that NMY-2 lost its cortical asymmetry in the NSM and identified two phosphorylation sites on NMY-2 that were partially dependent on PIG-1 for phosphorylation. They also found that phosphomimetic NMY-2 was able to partially rescue the loss of PIG-1 in the NSM division. An interesting set of future experiments would be to determine if PIG-1 has the same pattern of cortical asymmetry towards the larger side of other asymmetric divisions throughout *C. elegans* development, as we know from the difference between the localization of NMY-2 in the NSM neuroblast division and the Q lineage divisions that NMY-2’s localization pattern is not constant.

## Acknowledgments

Some nematode strains used in this work were provided by the Caenorhabditis Genetics Center, which is funded by the NIH Office of Research Infrastructure Programs (P40 OD010440). We thank Abby Dernburg, Chenshu Liu and Weston Stauffer for the use of their spinning disc confocal microscope and assistance in its use. We thank Nicolas Alexandre for his assistance with statistical analysis. and We thank Bob Horvitz and Guangshuo Ou for providing some of the strains used in this study. This work was supported by National Institutes of Health grant NS32057 to G.G.

## Supplementary Figures

**Fig S1:**
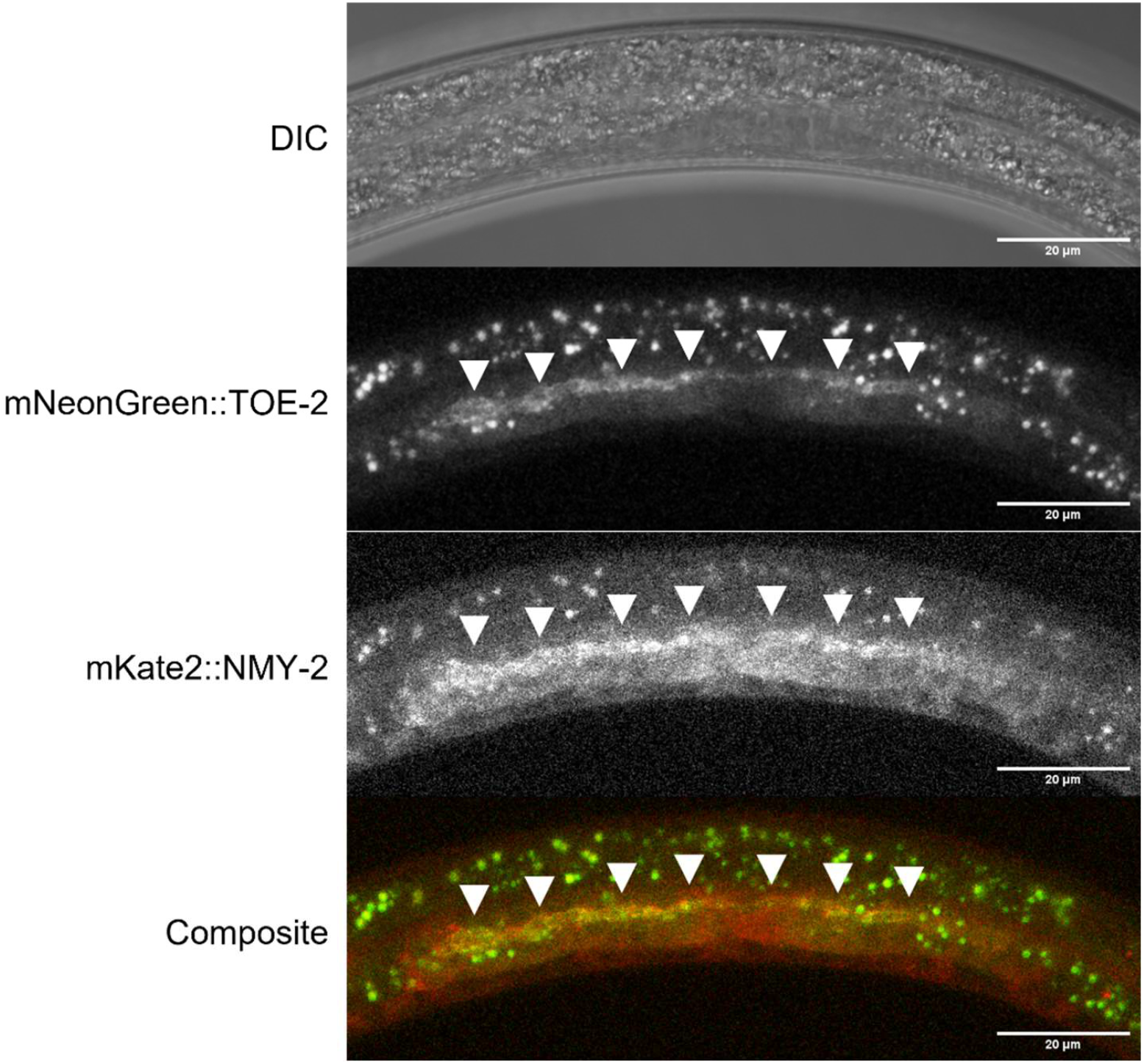
TOE-2 and NMY-2 are present at the apical germline. Confocal images of endogenously tagged TOE-2 and NMY-2 in a third larva stage (L3) hermaphrodite. Both proteins are expressed in the germline and accumulate at the apical surface of the germline cells. Arrowheads indicate the apical germline.

**Fig S2.**
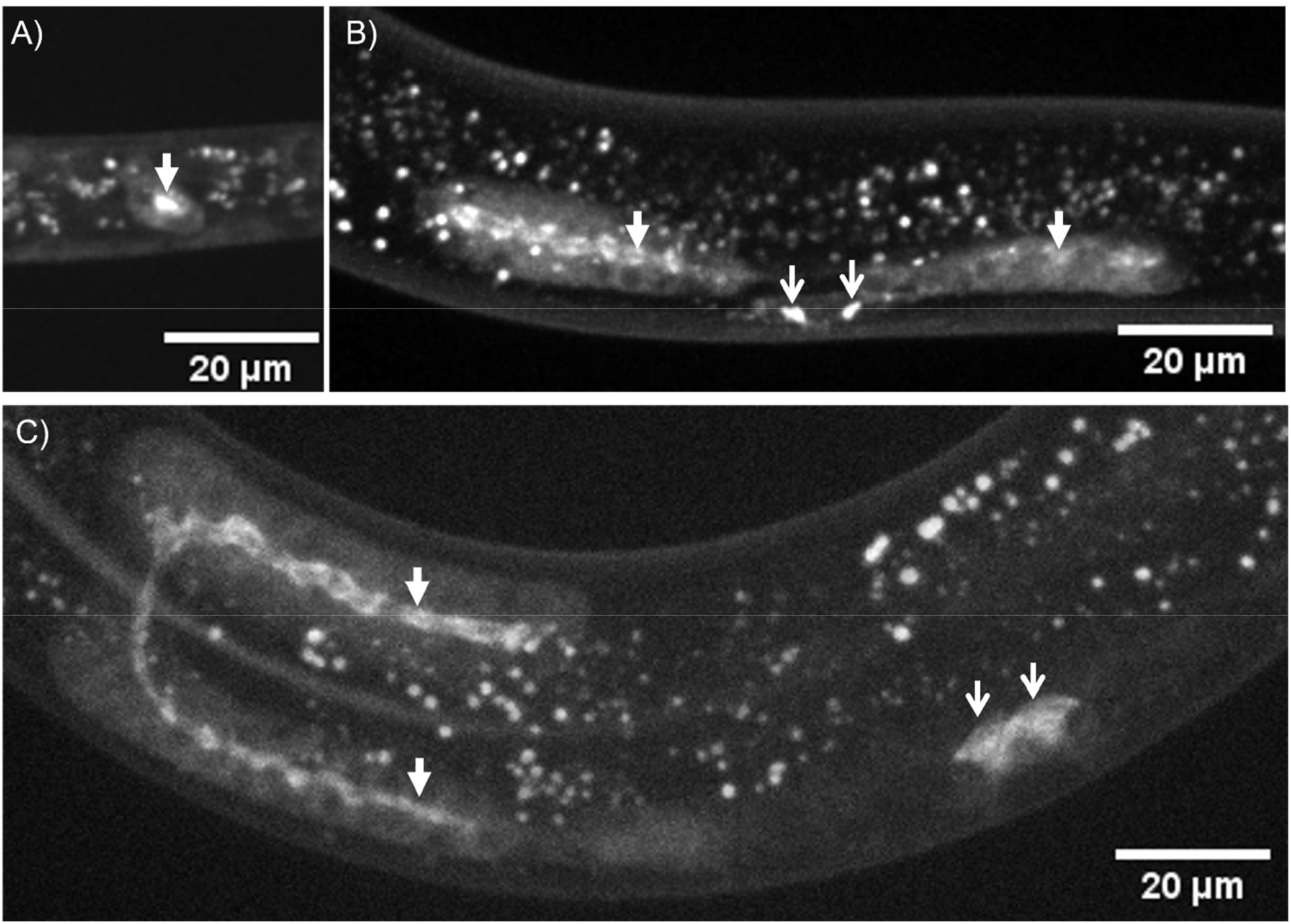
Confocal imaging of endogenously tagged TOE-2 localizing to the apical surface of the germ cells. A) mNeonGreen::TOE-2 in a first larval (L1) stage hermaphrodite. Closed arrowhead indicates mNeonGreen::TOE-2 at the point of contact between Z2 and Z3. B) mNeonGreen::TOE-2 in an L3 stage hermaphrodite. Closed arrowheads indicate localization of mNeonGreen::TOE-2 to the apical surface of the germline cells. Open arrowheads indicate mNeonGreen::TOE-2 localization to unknown cells near the vulva. C) mNeonGreen::TOE-2 in a fourth larval (L4) stage hermaphrodite. Closed arrowheads indicate mNeonGreen::TOE-2 localization to the apical surface of the germline cells. Open arrowheads indicate the positions of unknown cells near the vulva.

**Fig S3.**
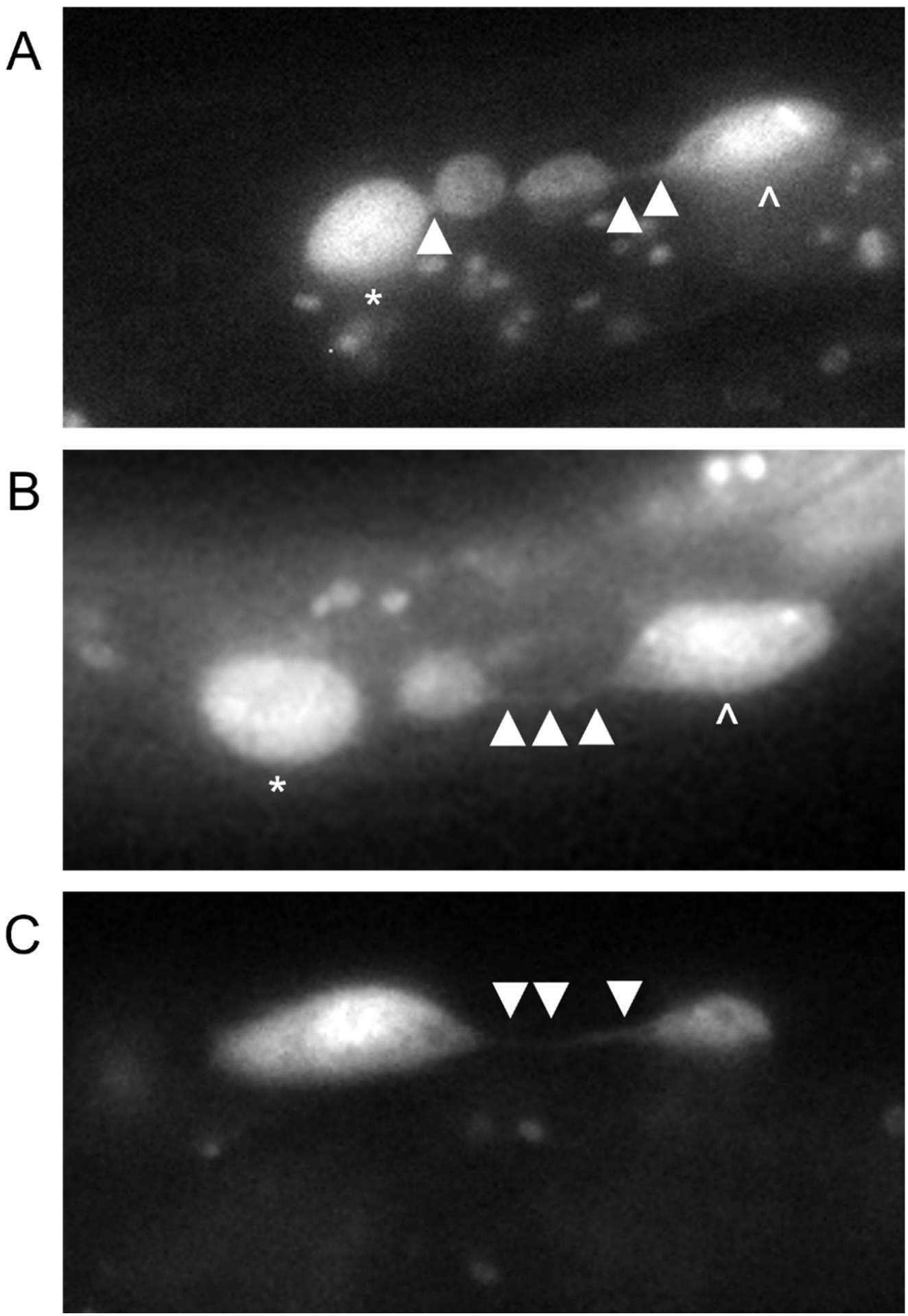
Intercellular bridges persist between Q lineage neuroblasts in *nmy-2(ts)* mutants. Anterior is to the left in A-C. A) QL.a and QL.p cells in an *nmy-2(ne3409ts)* mutant raised at the nonpermissive temperature with persistent intercellular bridges between their daughter cells. The cells on the left are the QL.p daughters. The QL.a daughters are more posterior because QL.a migrated past the Q.p cell before dividing. B) QL.a cell in an *nmy-2(ne3409ts)* mutant raised at the nonpermissive temperature with a persistent intercellular bridge between its daughter cells. The cell to the left is an undivided QL.p cell. C) QR.p cell in *nmy-2(ne1490ts)* mutant raised at the nonpermissive temperature with a persistent intercellular bridge between its daughter cells. Arrowheads indicate intercellular bridges, * indicates the QL.p and ^ indicates the QL.a cell.

